# Insulation of ribosomal promoter activity by Fis, H-NS or a divergent promoter within the packed *E. coli* genome

**DOI:** 10.1101/2024.03.27.586983

**Authors:** Elisa Brambilla, Malik Yousuf, Qing Zhang, Gladys Mbemba, Cyriane Oeuvray, Damel Mektepbayeva, Anne Olliver, Marco Cosentino Lagomarsino, Bianca Sclavi

**Author notes:** Authors contributions: BS, MCL: Formulation of theory and prediction BS, EB, AO: Contributions to experimental conception and design BS, EB, MY, QZ, CO, DM: Acquisition, analysis and/or interpretation of data BS, EB, MCL: Drafting the article or revising it critically for important intellectual content. Bianca Sclavi: UMR 7238 CNRS “Biologie Computationnelle et Quantitative” Sorbonne Université, Paris 75005 France. Marco Cosentino Lagomarsino: Dipartimento di Fisica, Università degli Studi di Milano, 20133 Milano and 14 INFN, Italy, and IFOM, FIRC Institute for Molecular Oncology, Milan, Italy. Elisa Brambilla: Department Microbial Population Biology Max Planck Institute for Evolutionary Biology. Qing Zhang: Beijing Institute of Genomics, CAS/China National Center for Bioinformation. Damel Mektepbayeva, Laboratory of biosensors and bioinstruments, National Laboratory Astana, Nazarbayev University, Astana, Kazakhstan 010000. The authors wish it to be known that, in their opinion, the first 3 authors should be regarded as joint First Authors.

## Abstract

Gene expression in bacterial cells is dependent on a gene’s position along the genome, mainly because of the effects of neighbouring genes’ expression, but also because of the local activity of nucleoid proteins, differing levels of DNA supercoiling and changes in gene copy number with growth rate and growth phase. This genome position dependence can be a source of specific regulation, however, in some cases it is necessary to have gene expression insulated from these local effects. *Escherichia coli* cells express ribosomal RNA from multiple operons found at different sites along the genome. The number of ribosomal operons varies in different strains and correlates with the maximal growth rate. rRNA promoters are under the regulation of Fis, H-NS, DNA supercoiling and ppGpp. These factors are known to result in growth phase and growth rate dependent regulation of gene expression. Here we show that the combined action of Fis and H-NS also provides insulation from the activity of both local and global regulatory factors. Furthermore, our results indicate that the presence of a divergently expressed gene can also act as an insulator revealing a DNA supercoiling gradient from the origin to the terminus. The organisation of ribosomal promoters therefore has been selected to allow for gene duplication independently of the influence of local genome organisation and neighbouring genes’ activity.

## INTRODUCTION

Bacterial genomes are very gene dense. Compared to eukaryotic genomes, they have relatively short stretches of non-coding DNA separating the genes. This can result in transcriptional interference by both run-on transcription by RNA polymerase (1, 2) and by the accumulation of positive and negative supercoiling downstream and upstream of the transcribing RNA polymerase (RNAP) respectively (3–5). The operon and supra-operon level of gene organisation is one way by which these effects can not only be minimized, but, in some cases, used to coordinate the expression of genes within the same pathway (6, 7). DNA topology and the binding of the abundant nucleoid-associated proteins have been proposed to act as delimiters of the evolutionarily conserved supra-operon level of organisation (6).

At a higher level of organisation, the distance from the replication origin of the genes coding for important growth-rate and growth phase regulatory factors is also found to be conserved (8, 9). One of the important factors influencing the rate of gene product formation is the change in gene copy number as a function of growth rate due to overlapping replication rounds (9–11). The binding of one of the major topoisomerase enzymes in *E. coli,* DNA gyrase, has also been known to increase towards the ori half of the genome (12). This can play a role in regulating the high levels of supercoiling created at the origin half of the genome by the increased rate of gene expression in exponential phase (13). Thus, while the rate at which DNA topology is created and regulated is likely to vary as a function of genomic position, the average level of supercoiling is found to be uniform along the genome in cells growing in exponential phase (14, 15). This suggests that there are at least two levels at which genomic position can influence the expression of a given gene: the local scale, influenced by neighbouring genes’ expression (3, 5), and at the genomic scale, depending on the distance from the origin (8), resulting in evolutionary constraints on gene position and orientation (16).

An additional level of regulation is dependent on the activity of the abundant nucleoid associated proteins (NAPs) that bind throughout the genome, with some preference for AT-rich DNA sequences (17–19). The NAPs have been shown to act like classical transcription factors either stabilizing or inhibiting the interaction of RNAP with a gene’s promoter sequence (20). However, the activity of nucleoid proteins can also influence gene expression by constraining DNA supercoiling and changing the local topology of DNA (21). The HU protein activity for example is correlated with gyrase binding and tightly linked with the amount of available superhelicity (22). Fis can act to activate transcription of stable RNA promoters when levels of negative supercoiling are low (23, 24). In the case of H-NS, the presence of higher affinity sites can nucleate the formation of extended binding domains that can silence transcription by stabilizing plectonemes and trapping RNA polymerase within loops (18, 25–27). NAP binding heterogeneity along the genome is thus an additional factor that contributes to the presence of clusters of co-regulated genes that can share membership to a common function or pathway (13, 28–30). NAP-rich regions are also associated with regions of the genome where transcription is silenced or decreased in a growth rate, growth phase and temperature-dependent fashion (26, 27, 31, 32).

Nucleoid associated proteins and DNA topology, together with RNAP availability, sigma factor competition and a set of small metabolites, such as ppGpp, have been identified as global regulators of gene expression because they can influence the activity of a large set of promoters, particularly those involved in growth rate and growth phase adaptation and general environmental stress response (33, 34). Here we have chosen to study the *rrnB*P1 promoter because it is a well-characterized promoter regulated by a set of global regulators, ppGpp, DNA topology, Fis and H-NS, and because it is found, albeit with some modifications, upstream of the seven copies of ribosomal RNA operons at different genomic positions (35–37).

In a previous work, we studied the gene expression levels from a reporter cassette where mut2-GFP expression is under control of the *rrnB*P1 promoter with a divergently expressed Kanamycin resistance gene inserted randomly in the genome by a transposon (38). We found that the insertion frequency was higher in AT-rich regions, but there was also a stronger repression of gene expression by H-NS at these sites.

Here we have inserted at several specific positions along the genome the same cassette containing the full-length *rrnB*P1 promoter and we compared its expression to a cassette containing a shortened version of the promoter lacking the Fis and the high affinity H-NS sites. Given the known role of Fis, H-NS and DNA supercoiling on the growth rate dependence of gene expression (21, 25), here we have studied the role of the upstream regulatory sequences at different growth rates, while our previous study was carried out in LB growth medium.

Previous studies have used a similar strategy in order to determine whether genome position could influence gene expression in *E. coli* as well as in other bacterial strains (5, 31, 32, 39–48). One of the main conclusions from these studies is that the results are strongly dependent on the choice of promoter used. Different promoters are reporters for different levels of regulation, such as transcription factor diffusion and availability (40), neighbouring gene transcription level and DNA supercoiling (5) or local nucleoid protein activity (31).

## MATERIALS AND METHODS

### Construction of the strains

The promoter-*gfp* KanR cassette construct fragment was amplified by PCR from the pDoc-K plasmid (49) and inserted at specific positions of the genome of the BW25113 *Escherichia coli* strain by the lambda Red recombination protocol (50). The kanamycin resistance cassette was removed by Flp-FRT recombination. The insertion positions were chosen to correspond to the position of the parS sites in the work by Espeli *et al*. (51) and are within intergenic regions, as defined by the gene positions in the Ecocyc Genome browser (52), where two genes converge, except for position Ori1 where the insertion is between two genes expressed in tandem (Table 1). The delta FIS and delta H-NS strains were created by P1 transduction into the KanR-less strains of the kanamycin resistance cassette inserted in either the *fis* or *h-ns* genes.

**Table 1.**
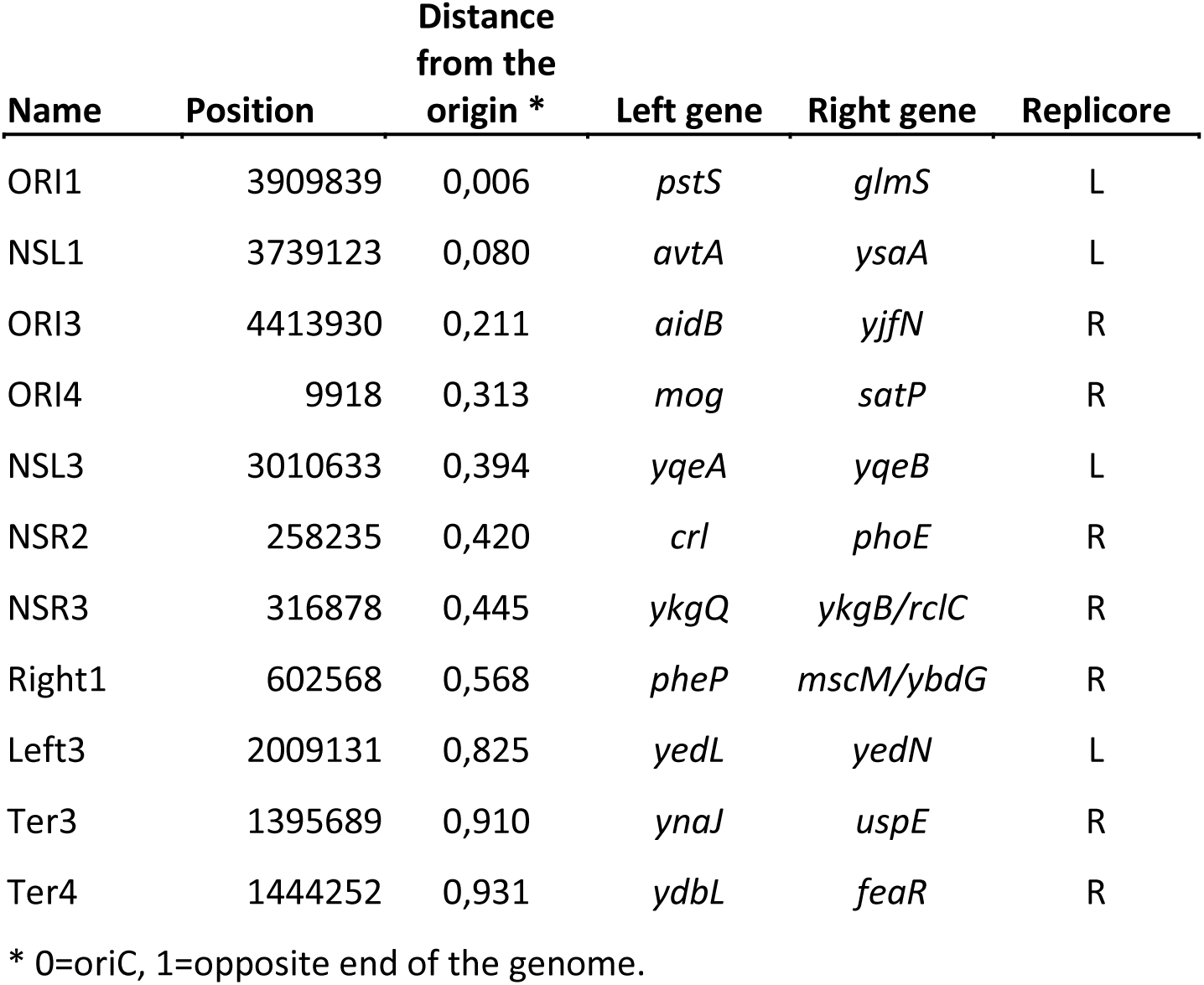
Position of inserted reporter cassette. . Each insertion site is an intergenic site where two genes converge, except for the ori1 site where the *pstS* gene is transcribed away from the *gfp* gene. The *mut2gfp* gene is transcribed towards the gene on the left, on the minus strand. The positions correspond to the Genebank entry U00096.2 sequence of the *E. coli* genome.

### Plate reader experiments and data analysis

Promoter activity and growth rate were measured as described in (31, 53) by growing the strains in a 96-well plate at 37°C in M9 minimal media supplemented with glucose (0.5 %), glycerol (0.5 %) and/or casamino acids (0.2 %) to vary the growth rate. The changes in optical density (OD_600_) and fluorescence were measured throughout the growth curve with a Tecan Infinite 200 PRO multimode reader. From these values, the growth rate (alpha: dOD/dt / OD) and the GFP production rate (promoter activity: dGFP/dt / OD) were measured (Figure 2). A window in time corresponding to mid- to late exponential growth phase was defined from which average growth rate and promoter activity were measured. The window was defined either by hand, by estimating the beginning and the end of the linear portion of the curve, or automatically from a linear fit of the semilog plot of the growth curve, giving very similar results. Changing the width of the window did not significantly affect the differences between the positions, indicating that the growth phase dependence of promoter activity is not influenced by the insertion position in a reproducible manner (data not shown). The normalization of gene expression for gene copy number as a function of growth rate and genome position was carried out by using the following equation:

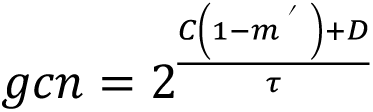

where *C* and *D* correspond to the duration of the C and D periods. The C period refers to the length of the process of DNA replication while the D period is the time between the termination of DNA replication and the end of cell division. *m’* defines the position of the gene with a value between 0 and 1, where 0 is the origin of DNA replication and 1 the polar opposite coordinate in the circular genome. The values of the C and D periods used for this analysis where obtained from Stokke et al. (2012) where they studied *E. coli* DNA replication in very similar experimental conditions to those used here (54). For the Gyrase inhibition experiments Novobiocin at 50 µg/ml final concentration was added to the cultures in the 96-well plate once the cells had reached early exponential phase (OD_610nm_ 0.02 in the plate reader).

## RESULTS

In order to better understand the effect of a promoter’s genome position on the activity of transcription regulatory factors such as Fis, H-NS and DNA topology, we have chosen to compare the activities of the following promoters: the full-length *rrnB*P1 promoter (P1long); a shortened version lacking the three Fis sites and one of the three H-NS sites (P1short) (Figure 1A) and a constitutive promoter from T5-phage that has a consensus −10 and −35 sequences separated by a 17 bp spacer and an AT-rich UP element (P5 promoter) (SFig. 1). To measure promoter activity we cloned these promoters upstream of the *mut2-gfp* gene and inserted this construction, including a divergently oriented kanamycin resistance cassette (KanR cassette), in several sites along the genome (Figure 1B, Table 1 and SFig. 1). There are no terminator sequences downstream of the two genes. The naming of the insertion sites correspond to the insertion positions used by Espeli et al (51) and therefore are related to the name of the macrodomain where they are found, *ori*, non-structured left and right (*nsl*, *nsr*), *left*, *right* and *ter*. An example of the data obtained by the plate reader is shown in Figure2A.

**Figure 1.**
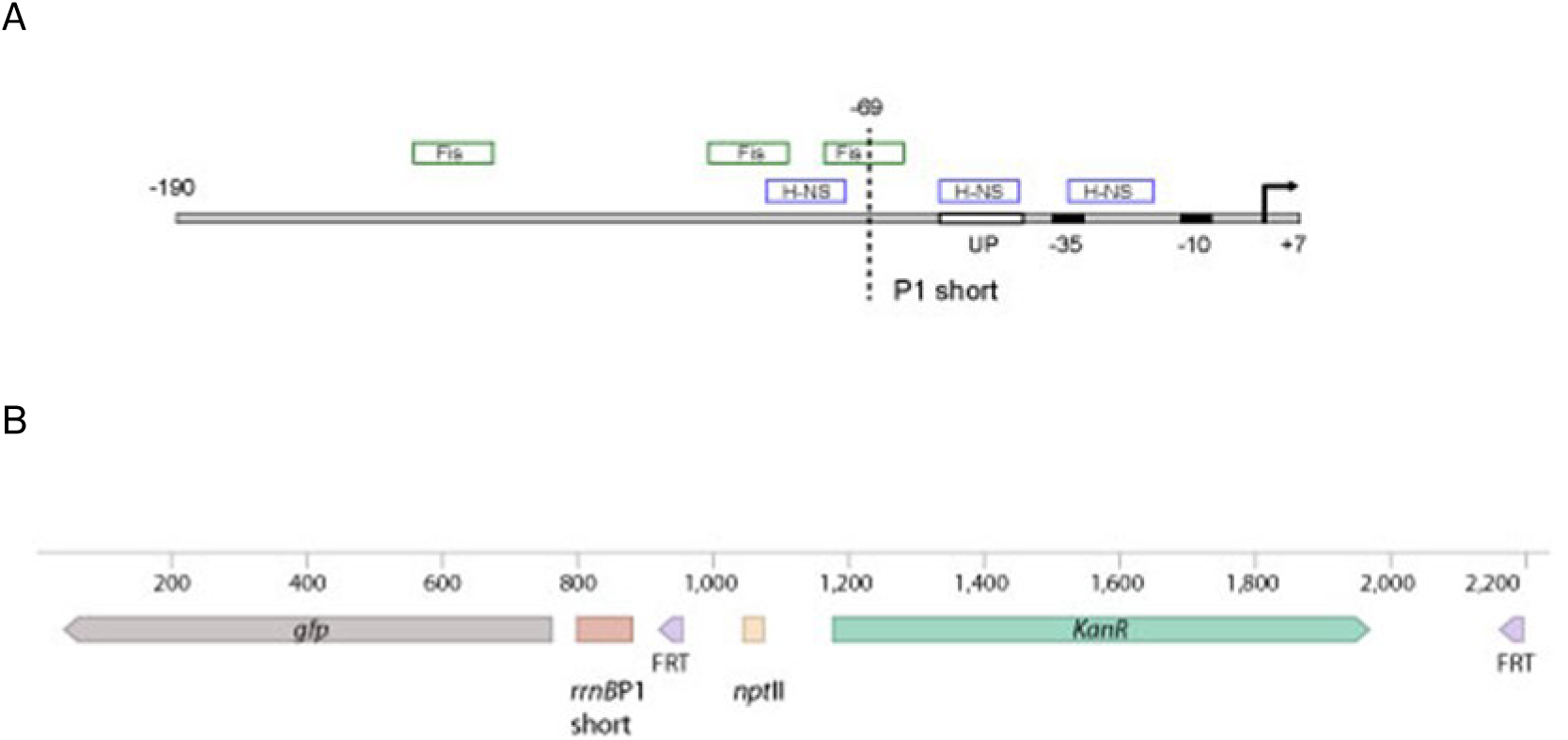
Promoter-reporter construct. A. The *rrnB*P1 promoter, in its full-length, P1long, and shortened P1short versions, was placed upstream of the *mut2-gfp* gene with a strong RBS (B). The divergently expressed KanR cassette is transcribed from its original *npt*II promoter (55) and is flanked by two FRT sites that allow for excision of the cassette by Flp recombinase.

**Figure 2.**
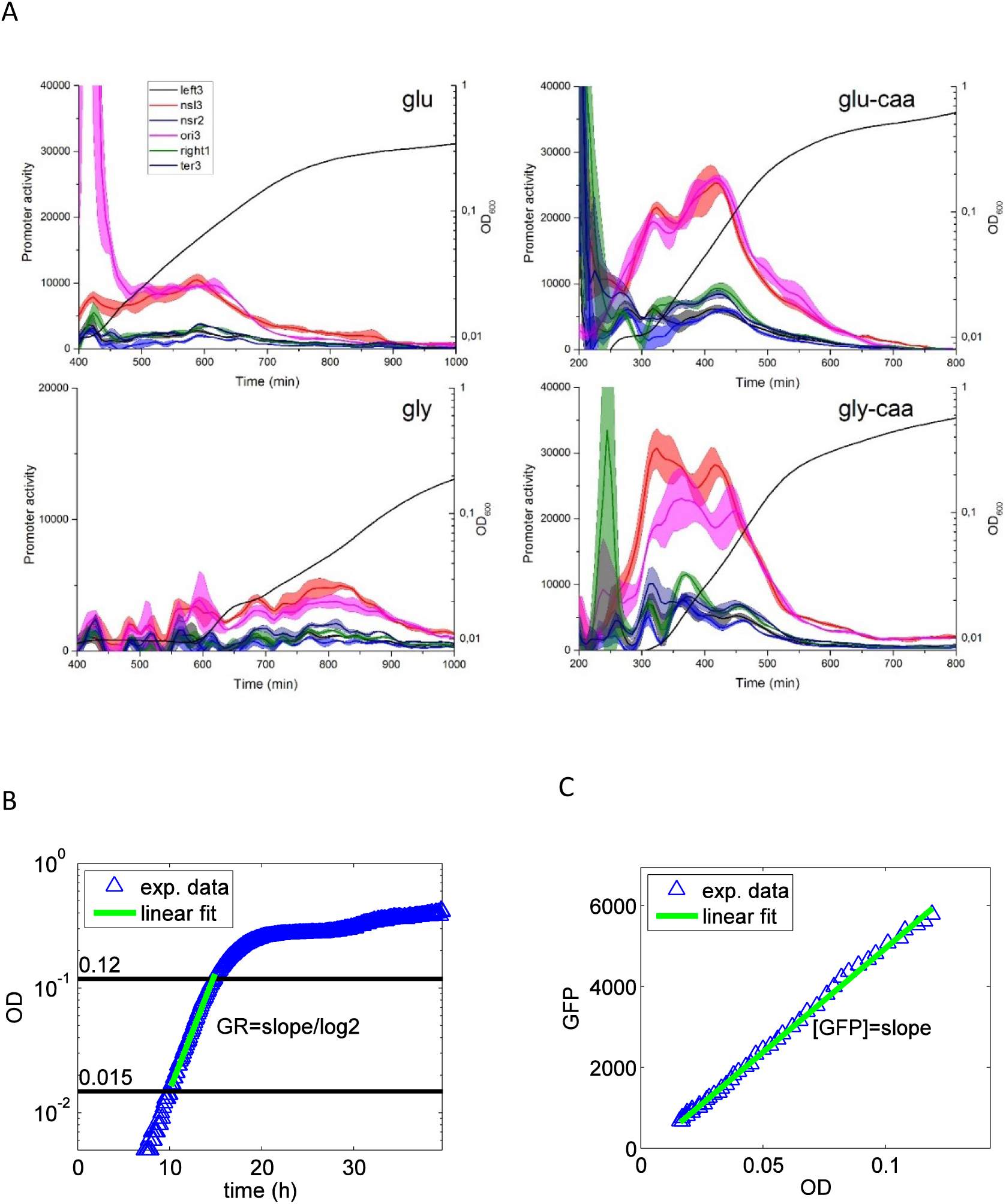
**A. Promoter activity (dGFP/dt / OD) for P1short without the KanR cassette in six different positions and in four growth media.** The four growth media labels are based on M9 minimal medium and defined as: glu: glucose (0.5 %), gly: glycerol (0.5 %), glu-caa: glucose (0.5%) and casamino acids (0.2 %), gly-caa: glycerol (0.5%) and casamino acids (0.2 %). The change in OD_600_ is shown by the black line without shading. The coloured shading around each line corresponds to the error bar (SEM) is from the three replicates within the same plate. Each colour corresponds to a different insertion position. Variations in the low levels of OD in early exponential phase cause the appearance of non-reproducible peaks in the derivative of fluorescence normalized by OD. Data for the other promoter constructs can be found in SFig. 2. **B.C. Automated data analysis protocol.** Growth rate (GR), GFP concentration and GFP production rate for all the strains were derived from the data of the optical density at 600 nm (OD) and green fluorescence (GFP) as a function of time (labelled as exp.data). The exponential phase was defined as the period during which log(OD) is linear and was defined by two thresholds (the thresholds were set manually or by an automatic method (56, 57) both of which give similar results). Based on the data in exponential phase, the growth rate was defined as the slope from the linear fit of the plot of log(OD) versus time divided by the log of 2. GFP concentration was derived from the slope of the linear fit of the plot of GFP versus OD, and the GFP production rate was the product of GFP concentration and growth rate.

### A divergent promoter-gene cassette can increase rrnBP1 promoter activity in a growth rate and genome position dependent manner

In order to measure promoter activity independently of gene copy number effects, we normalized the promoter activity values by the gene copy number expected for that genome position at the growth rate of the sample (see Materials and Methods and (31)). The results obtained with the P1short promoter show that there are significant differences in promoter activity depending on insertion position (Figure 3), as observed previously for the *lac* and TetO1 promoters (5, 32, 48). We decided to compare the activity of the promoter in the presence and absence of the divergently oriented KanR cassette mainly because transcription of the *Kan* gene can create negative supercoiling upstream of the RNA polymerase (RNAP) and increase the expression of the *rrnB*P1 promoter, known for its sensitivity to the levels of negative supercoiling (58). Since strong, constitutive promoters such as P5 are known to be less sensitive to changes in negative supercoiling than the stable RNA promoters containing a GC-rich discriminator region (59, 60), the comparison of the effects of the KanR cassette on the P1 and P5 promoters can help distinguish between the increase in GFP expression due to the change in the local levels of negative supercoiling versus other possible effects of the presence of the KanR cassette.

**Figure 3.**
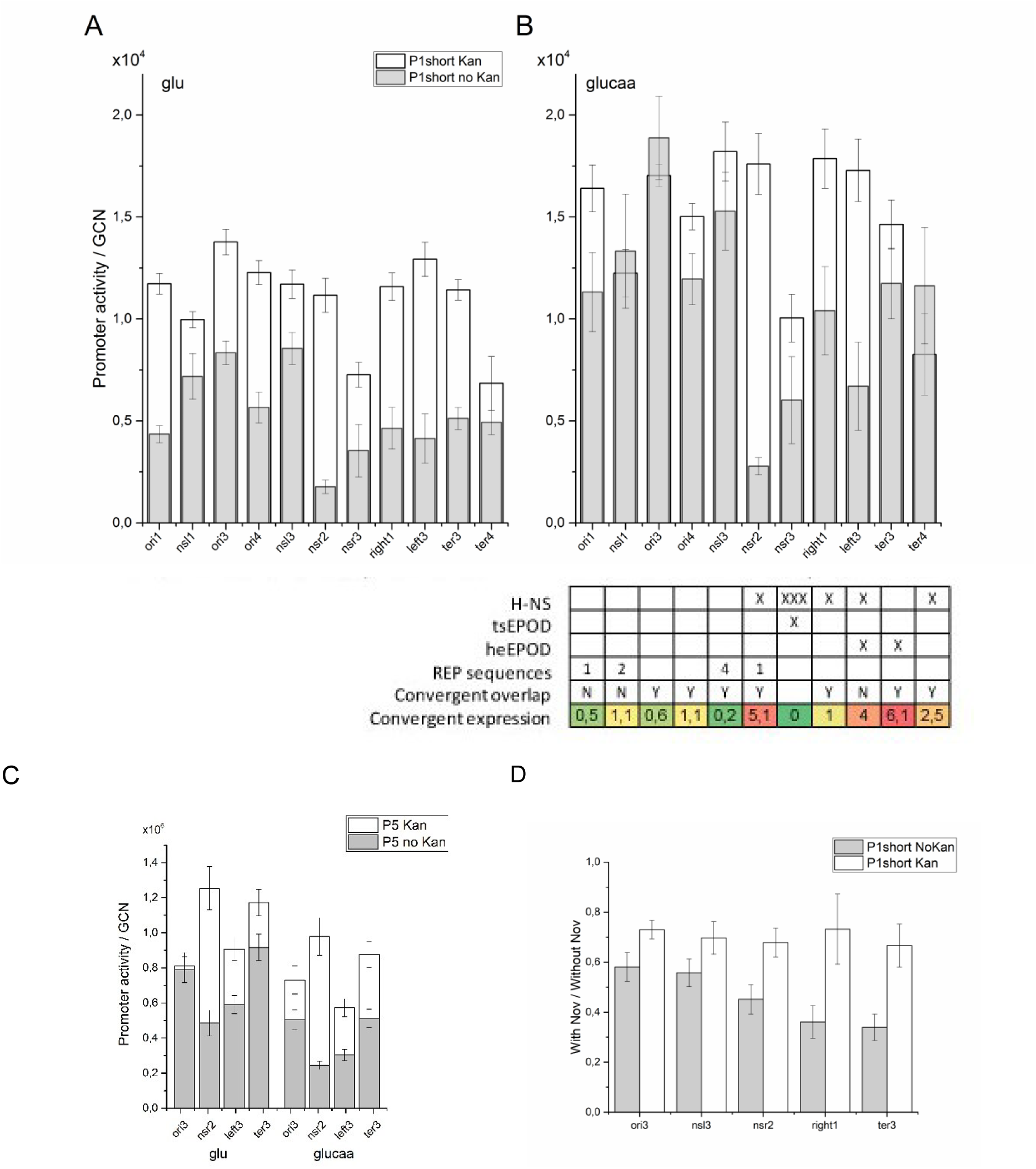
Activation of promoter activity by a divergently expressed gene as a function of genome position. Average promoter activity in exponential phase at different insertion sites along the genome was measured by the change in fluorescence as a function of time normalized by OD_600_. The sites are ordered from left to right as a function of their distance from the origin. **A.B.** *rrnB*P1 promoter activity is shown in the presence and absence of the divergently expressed KanR cassette in two growth media, M9-glu (**A**) and M9-glucaa (**B**). The table summarizes data from previous studies at each position: H-NS occupancy (18), X corresponding to binding regions of about 1 Kb and XXX a larger region of about 15 Kb, ts and heEPOD (26), the presence of rep sequences in the intragenic region, the possible overlap of transcription from the convergently expressed gene (1) and the level of expression of the convergent gene (61). The colour, from green to red, corresponds to the level of expression. **C.** Gene expression with and without the divergently expressed KanR cassette for the P5 promoter, in M9-glu and M9-glucaa. **D.** Ratio of gene expression with and without novobicin (50 µg/ml final concentration) in the presence and absence of the KanR cassette in M9-glu. Promoter activity is measured in arbitrary units determined by the acquisition settings of the plate reader. The same acquisition settings were used for the experiments shown in the four panels. The error bar is the SEM from three independent experiments.

In the absence of the KanR cassette, a significant decrease in promoter activity is observed at most insertion sites for both promoters (Figure 3). However, compared to the P5 promoter, for P1short this decrease is greater at the slower growth rate, in M9-glucose (M9-glu), than in M9-glucose-casamino acids (M9-glucaa) (SFig. 3). This is consistent with a lower level of negative supercoiling in the cells growing at a slower growth rate and thus a stronger effect of the presence of a divergently transcribed gene.

To confirm that the effect of KanR on P1 can take place via changes in negative supercoiling we have measured promoter activity after the addition of sublethal levels of the gyrase inhibitor novobiocin in the M9-glu growth medium (Figure 3D). Limitation of gyrase activity resulted in a stronger decrease in the activity of the P1short promoter in the absence of the KanR cassette than in its presence. Interestingly, while the decrease in promoter activity in the presence of the KanR cassette is around 30% for the five insertion sites studied here, in the absence of the KanR cassette, the decrease depends on the distance from the origin: the insertions at ori3 and nsl3 decrease around 40%, while nsr2, right1 and ter3 decrease up to 70% compared to the level of expression in the absence of the drug. The presence of the divergent gene thus buffers the reporter cassette promoter from variations in supercoiling along the genome. Furthermore, this result supports the trend that is already visible in the absence of novobiocin, where the activation by the divergently expressed gene tends to be greater towards the terminus end of the genome. Finally, we have carried out the control where the kanamycin resistance cassette has been reversed, to be transcribed towards the promoter, instead of away from it. This results in a decreased level of expression, similar to the one observed in the absence of KanR, showing that the orientation of the upstream gene is important for activation, consistent with a DNA supercoiling dependent effect (SFig. 7).

The effect of the divergent KanR cassette depends on the insertion position. These differences are enhanced at the faster growth rate when one compares the sites at the *ori* and *nsl* macrodomains to those farther from the origin.

Figure 3 includes a table that summarizes the available information on the different factors that could influence gene expression at a given position: the presence of H-NS rich regions (18), tsEPODs (transcriptionally silent Extended Protein Occupancy Domains) and heEPODs (highly expressed Extended Protein Occupancy Domains) (26), the presence of rep sequences in the intragenic region, as possible targets of gyrase activity (62), the possible overlap of transcription from the convergently expressed gene (1) and the level of expression of the convergent gene (61). The level of expression of both genes, on either side of the insertion, is shown in SFig. 4. More details about the larger (40 Kb) genomic context of the insertions can be found in SFig. 5.

The effect of the KanR cassette on P1short expression at the *nsr2* and *left3* positions remains strong independently of growth rate (SFig. 3). At both positions, there is a highly expressed gene convergent to the reporter gene and a low level of transcription coming from upstream due to H-NS binding. The main difference between these two positions is that the *yedL* gene at the *left3* position has a strong terminator downstream of the insertion site, while the *crl* gene at the *nsr2* position has a weak terminator at the end of the gene and a stronger one 217 base pairs downstream, past the insertion site of the reporter cassette (1). Therefore, in the latter case it is probable that transcription interference contributes to the low level of expression of GFP at the *nsr2* position when the promoter is weaker in the absence of the KanR cassette. The different magnitude of the effects of the KanR cassette observed for the activity of P1short (6.5-fold) and P5 (3-4 fold depending on the growth rate) at the *nsr2* position can be explained by the greater susceptibility of P1short to the changes in the local DNA topology due to the positive supercoiling created by the convergent gene’s transcription towards the *mut2gfp* gene. A similar result is observed to a lesser extent at the *left3* position (SFig. 3). The *ter3* position also has a highly expressed gene convergent to the reporter gene; however, in this case, there is no H-NS binding upstream instead there is the highly expressed *uspE* gene.

Transcription from neighbouring genes can change the local levels of negative supercoiling (3). All the insertion sites except for *ori1* are found between two convergent genes; therefore, we expect that the local levels of positive supercoiling will increase with the level of expression of neighbouring genes. We used data from gene expression levels in M9-glucaa (61) to estimate the levels of transcription-coupled DNA supercoiling (TCDS) by the method of Sobetzko (3) from the transcription occurring over a 5Kb window on either side of the insertion site (SFig. 4). However, we did not find a strong correlation between TCDS and promoter activity, especially at the *ter3* position, where high expression of neighbouring genes on either side would predict a high level of positive supercoiling. The lack of a correlation might be explained by the fact that all insertions are within the same kind of local gene orientation, in addition to the effect of the presence of nucleoid proteins binding regions, such as H-NS (see Discussion).

Our previous work had shown that the insertion of an H-NS repressed promoter within an extended H-NS binding domain resulted in increased H-NS repression compared to the same promoter inserted at other sites on the genome (31, 38). The P1short promoter still contains two out of the three H-NS sites found in the full-length *rrnB*P1 promoter, albeit the missing site is the only one that does not directly compete with RNAP for DNA binding, and its absence has been shown to weaken significantly repression by H-NS (63). In this dataset, the *nsr2*, *right1*, *left3* and *ter4* sites all are within H-NS binding domains of about 1 Kb (18) (SFig. 5). The *nsr3* site is found within an extended H-NS binding domain (about 15 Kb), and is the only site that is found within a transcriptionally silent extended protein occupancy domain (tsEPOD) (26). At the faster growth rate, while at *nsr2*, *right1* and *left3* the presence of the KanR cassette has a strong effect on promoter activity, at *nsr3* promoter activity remains lower than the others even in the presence of the KanR cassette, indicating that the additional 1.2 Kb added by the KanR cassette, while decreasing repression by H-NS, is still not sufficient to disrupt the effect of the extended H-NS binding domain.

The *right1* and *ter4* sites are within very similar local contexts, but their level of expression and the effect of the KanR cassette are quite different. This is consistent with previous results showing that repression by H-NS tends to be stronger at sites closer to the terminus of replication (31).

To obtain a more direct measure of the effect of H-NS on gene expression, the chromosomal insertions lacking the divergent KanR cassette, where H-NS repression in stronger, were studied in cells where H-NS expression was abolished by replacing the *hns* gene with an antibiotic resistance cassette (Figure 5A). GFP expression from the cassettes within H-NS rich regions increased to reach a similar level of the other sites, independently of the growth rate.

### Fis activation of the rrnBP1 promoter depends on the insertion position and renders it less sensitive to local context

The full-length *rrnB*P1 promoter, P1long, contains three Fis sites and an additional H-NS site upstream of the promoter. The comparison between P1long and P1short allows us to gain an improved understanding of the role of these two proteins as a function of genome position. The concentration of Fis and H-NS and the cellular levels of available negative supercoiling are known to depend on growth rate (20, 21, 64), we thus studied promoter activity in four different growth media, M9-glycerol (0.5 %), M9-glucose (0.5 %), M9-glycerol (0.5 %) and casamino acids (0.2 %) and M9-glucose (0.5 %) and casamino acids (0.2 %). Because of the strong dependence of *rrnB*P1 activity on growth rate, we compared GFP expression in the strains containing P1long and P1short in the same experimental conditions (Figure 4). The results obtained confirm the ones shown in Figure 3 and clearly show the growth rate dependent activity of both the P1short and P1long promoters (36) independently of genome position and gene copy number (SFig. 6). The growth rate dependence of the shortened version of the *rrnB*P1 promoter activity, in the absence of the Fis sites and the full set of H-NS sites, is known to be mainly due to the changes in the levels of ppGpp and DNA supercoiling (21, 36, 65–67). As expected, the samples in the two growth media with casamino acids, and thus low levels of ppGpp, show a similar level of promoter activity (66).

**Figure 4.**
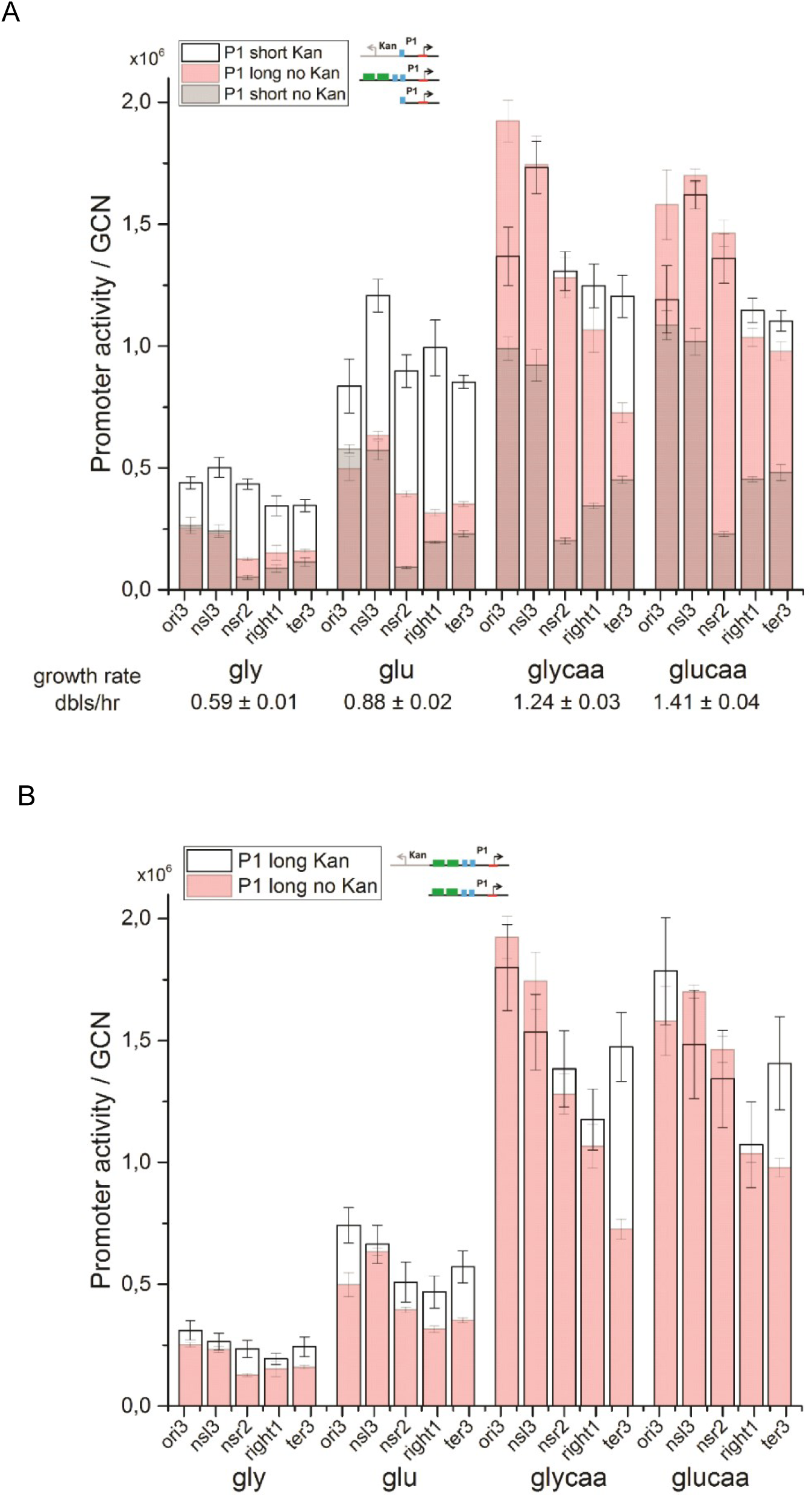
**A. *rrnB*P1 promoter activation by Fis or by a divergently expressed gene as a function of growth rate.** Promoter activity normalized by gene copy number at five positions in four different growth media. The resulting growth rates, averaged over all the strains, are shown underneath each set. Comparison of the *rrnB*P1 short promoter, in the presence and absence of the divergent kanamycin resistance cassette and the full-length *rrnB*P1 promoter in the absence of the divergent kanamycin cassette. **B**. **Comparison of the *rrnB*P1 long promoter, in the presence and absence of the divergent kanamycin resistance cassette.** Promoter activity is measured in arbitrary units. The error bar is the SEM from three independent experiments. The error for the growth rate value is from all the samples in that given growth condition.

The addition of the Fis sites and the presence of the full set of H-NS binding sites changes the growth rate dependence of promoter activity, this is due mainly to an increase at the faster growth rates, in accordance with the known growth rate dependence of Fis expression and the increased repression of ribosomal promoters by H-NS at slower growth rates (68, 69). In the presence of casamino acids, comparison of P1long without KanR and P1short with the KanR cassette shows that they both have an increased promoter activity compared to P1short. The extent of activation by Fis is of the same order of magnitude than what was previously described (70). On the other hand, at the slower growth rates the KanR cassette activates the *rrnB*P1 promoter more than Fis, whose effect is minimal. The KanR cassette is transcribed by a weak but constitutive promoter (55), whose activity is therefore mostly growth rate independent, while Fis activity increases with increasing growth rate and at slower growth rates it competes with repression by H-NS.

At the faster growth rates, it is interesting to note the similarity in the extent of activation as a function of genome position between the effect of the divergent KanR cassette and the Fis sites, indicating that Fis plays a similar role to the divergent gene in the activation of the promoter and thus effectively screening it from the effects of transcription at neighbouring genes and the binding of H-NS, this is particularly striking at the *nsr2* position.

The results obtained in the strains lacking Fis support this interpretation (Figure 5B). The lack of Fis in the cell is known to increase the levels of negative supercoiling (71), indeed expression from the P1short promoter increases in this case, especially at the slower growth rate, in M9-glu, where the levels of negative supercoiling are lower to begin with in the wild type context. At the faster growth rate, the two sites closest to the origin are the least affected, consistent with a higher level of negative supercoiling at the ori end of the genome. The H-NS dependent pattern of site-specific repression however is retained. The lack of Fis has the opposite effect on the P1 long promoter, the loss of activation is greater than the effect of increased negative supercoiling, particularly at the faster growth rate (Figure 5C). In the M9-glu growth medium, increased repression by H-NS at the nsr2 site now becomes apparent, underlining the role of Fis in contrasting the activity of H-NS.

**Figure 5.**
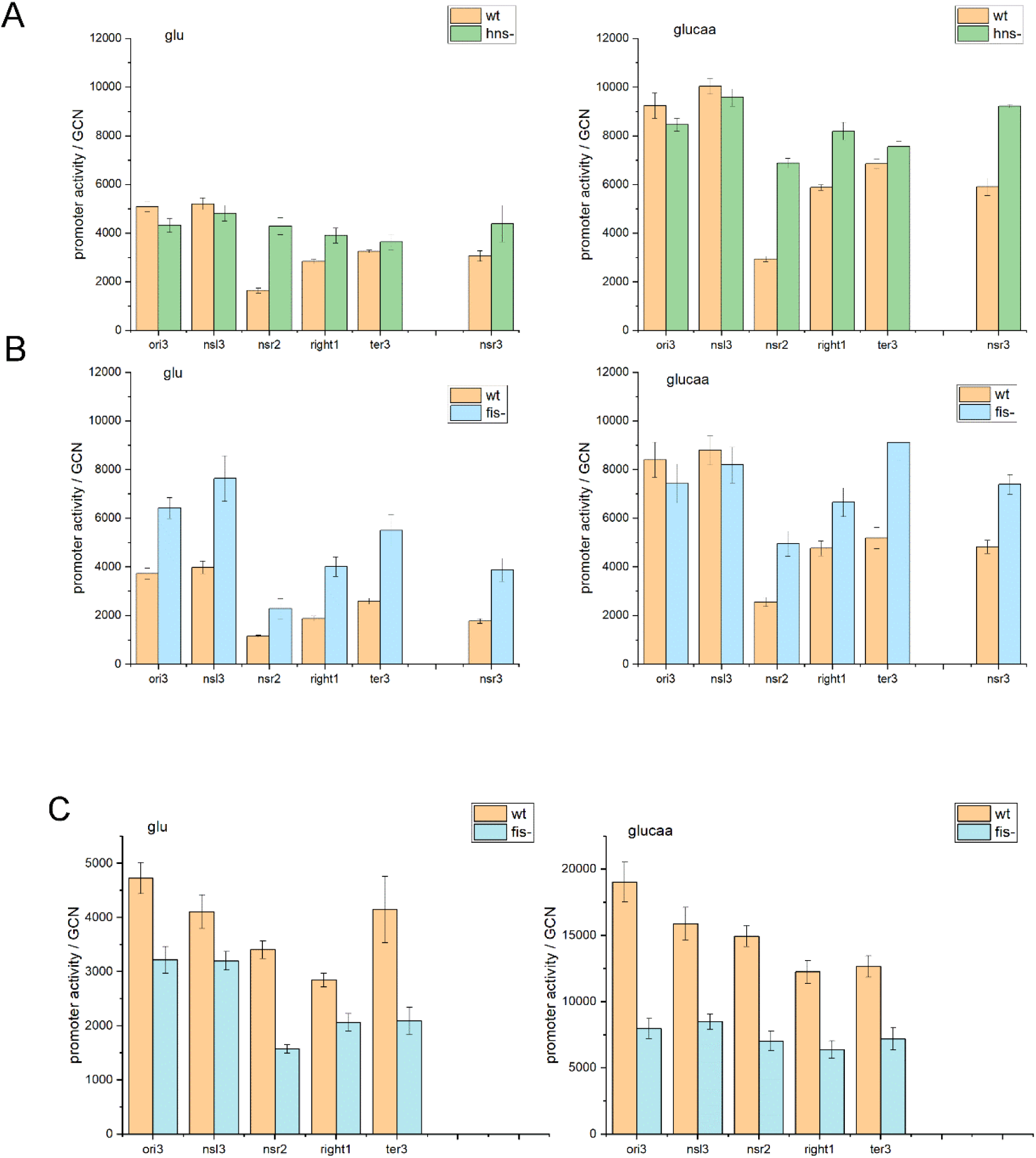
Expression of mut2-GFP from the P1short (A and B) or P1long (C) in strains lacking either H-NS (A) or Fis (B and C). Promoter activity is measured in arbitrary units. The error bar is the SEM from three independent experiments. The error for the growth rate value is from all the samples in that given growth condition.

In contrast to what is observed at the P1short promoter, the presence of the KanR cassette upstream of the P1long promoter does not have a very strong effect on promoter activity, independently of the growth rate (Figure 4B). Activation by Fis, mainly at fast growth, and repression by H-NS at slow growth can limit the effect of this neighbouring divergent gene on the activity of *rrnB*P1.

In most cases, the expression of the GFP reporter gene does not seem to influence significantly the growth rate of the cell. When a decrease in growth rate is observed, it is in the richer growth media for the strains containing the reporter cassette at the two positions closest to the origin, *ori3* and *nsl3*, and thus when gene dosage is increased at the same time as promoter activity (SFig. 6). This is consistent with previous studies showing that the expression of the heterologous GFP protein can decrease the amount of available resources required for growth (72). One specific condition however causes a large decrease in growth rate: the full-length *rrnB*P1 promoter in the absence of the KanR cassette at the *ter3* position. This position is within a highly expressed Protein Occupancy Domain (heEOPD) (26) and specifically between two convergent genes, *ynaJ* and *uspE,* that are among the most highly expressed compared to the neighbours found at the other insertion sites. *ynaJ* codes for a predicted inner membrane protein while *uspE* codes for a stress-response protein involved in cell motility (52). It is interesting to note that in the presence of the divergent KanR cassette this effect on the growth rate of the cell is not observed, despite a greater amount of GFP being produced. The expression of the KanR gene not only insulates the expression of the reporter protein, but can also act to insulate the neighbouring genes from the inserted cassette, as it has been shown recently for Fis binding (24).

## DISCUSSION

Bacterial genomes evolve by both intragenomic and intergenomic, horizontal, gene transfer (73). Gene expression can be influenced by its position in the genome due to both local and genomic scale effects (5, 8, 31, 42, 43). Therefore, when a gene lands in a new position in the genome it will not necessarily be expressed in the same fashion as before (31, 38). In some cases, this can be an advantage and the new position will be maintained, such as piggy-backing on already existing local regulatory elements. In other cases it is preferable to keep at least some of the same levels of regulation as before the transfer.

One example of intragenomic gene transfer is the duplication of ribosomal operons that is associated with strains that can rapidly adapt to faster growth rates (74). Here we find that regulation of a ribosomal promoter by Fis and H-NS decreases the variability in promoter activity between the different insertion positions (from ∼5 to ∼1.3 fold), thus maintaining the resulting growth-rate-dependent expression independent of the activity of neighbouring genes. This type of insulation by both a repressor and an activator can insure that the response remains proportional to the input, in this case growth rate, at both low and high levels of promoter activity (75). The main influence of the gene’s position will then remain the change in gene copy number, which can result in an increase in the average ori to ter ratio of up to 2.5 times during growth in rich medium.

The number of ribosomal promoters in the genome varies and is proportional to the maximal growth rate that can be attained by the cell (74). Each of the copies of ribosomal promoters is activated by two to five Fis binding sites and is activated by a different extent when measured independently of its genomic context (37, 70, 76). Not all seven ribosomal operons are found within the same kind of local context, in some cases the upstream gene is expressed divergently to the ribosomal promoter. The genomic context of the native *rrnB*P1 promoter however shows that the two upstream Fis sites are found within the coding region of the essential *murI* gene, which is expressed co-directionally with the *rrnB* operon (52). Other stable RNA promoters are regulated by a very similar promoter organisation. At the *tyrT* promoter region, for example, one of the three Fis sites is also found within the reading frame of the *purU* gene transcribed co-directionally. The presence of H-NS binding sites and a variable number of Fis sites upstream of the promoter region could facilitate gene duplication by maintaining the growth rate dependence of gene expression independently of its specific location in the genome and the possible changes in the gene expression level of neighbouring genes.

### Supercoiling, local vs global (growth rate) effects

The factors that have been observed to influence the local genomic context include the local concentration of nucleoid proteins (31) and the orientation and the level of expression of neighbouring genes resulting in transcriptional interference and changes in the local levels of available DNA supercoiling (1, 3, 5). Transcription of a divergent gene, such as the KanR cassette, will change the DNA topology upstream of the RNA polymerase resulting in an increase of the local levels of negative supercoiling (4). At the *rrnB*P1 promoter, a change in DNA topology is expected to have a stronger effect than at the P5 promoter since it contains a GC-rich discriminator region (59, 60). An effect of the presence of the KanR cassette is observed for both promoters, in part due to the influence that negative supercoiling can have on the RNAP elongation rate (77). The difference between the effects of the KanR cassette on these two promoters can therefore be an indication of the changes in negative supercoiling felt at the promoter. Transcription from the KanR cassette promoter is not very strong (55), but it suffices to increase local levels of negative supercoiling and activate the P1 promoter. In addition, the presence of the KanR cassette also increases the distance from the promoter to the genes that may be expressed upstream by about 1.2 Kb, decreasing the effect of the positive supercoiling resulting from convergent gene expression (SFig 4C).

A greater increase in P1short promoter activity by the presence of KanR compared to P5 is observed both as a function of insertion position and as a function of growth rate. The former reflects the changes in the local level of supercoiling and thus of the transcription of neighbouring genes, the latter is observed as a generalized increase in the effect of KanR for P1short at all positions at the slower growth rates. This is in agreement with the idea that the average levels of available negative supercoiling in the genome decrease with decreasing overall transcription activity and thus growth rate (15, 21). The levels of ppGpp also increase in the growth media lacking casamino acids (66). Increased local negative supercoiling by KanR can act to counter the negative effect of ppGpp (21). This aspect can be addressed in the future for example by using a promoter that is sensitive to supercoiling but not ppGpp levels, such as a ribosomal promoter sequence containing a mutation at −7 that renders it less sensitive to ppGpp but maintains the properties of the discriminator region (78).

DNA gyrase sites can also act as insulators of changes in supercoiling due to neighbouring genes’ activities (79). In this case, the difference in the rates of local supercoiled DNA production by transcription and DNA replication and the rate at which gyrase can act on the DNA will determine the net amount of available supercoiling (77). The data on the P1short promoter and on the effect of the KanR cassette upon gyrase inhibition are suggestive of a gradient of negative supercoiling from ori to ter. Previous experiments using psoralen crosslinking had observed an HU-dependent increase of negative supercoiling at the terminus in stationary phase cells, but a lack of a gradient in exponentially growing cells (14). Promoter activity might be a more sensitive reporter of the levels of negative supercoiling than the ones measured by crosslinking, notably crosslinking averages over a time window of several minutes and is thus unable to detect fluctuations in the conformation.

### The role of Fis

The role of Fis in the activation of ribosomal promoter activity has been studied mainly outside the context of the promoter’s genomic position. The early studies *in vitro* and on plasmids *in vivo* have played a crucial role to define the effect of Fis bound to the sites upstream of the *rrnB*P1 promoter and other closely related stable RNA promoters such as *tyrT* and *rrnA*P1 (59, 70, 80–82). These studies have shown how a direct interaction between Fis and the alpha subunit of RNAP can stabilize the transcription initiation complex (81, 82). In addition, the specific binding of Fis upstream of the promoter can wrap the DNA, change the local twist to destabilize the GC-rich discriminator region and favour promoter melting, acting as a “topological homeostat” (80). Activation by Fis was thus shown to be more significant in conditions where the levels of negative supercoiling were suboptimal for promoter melting (59). Finally Fis binding can also result in a supercoiling diffusion barrier (24, 80). Here we show that activation by Fis can vary from hardly any effect to a 6-fold effect (Figure 4) depending on genomic position, showing to which extent the local genomic context can influence the study of the activity of promoters *in vivo*. The degree of activation by Fis can depend on the local levels of negative supercoiling and on the competition with H-NS binding, depending on the local activity of H-NS.

It is interesting to note the similar magnitude in the activation of P1short by the addition of the KanR cassette and the Fis sites at the faster growth rates. The KanR cassette therefore also results in a decrease in the variability of promoter activity as a function of genomic position. However, unlike Fis, the expression of the KanR is not growth rate dependent; therefore, it cannot provide the fold-change in expression required for the adaptation of ribosomal promoters to different growth rates (SFig. 6). Fis and the KanR cassette activate the P1 promoter by seemingly different mechanisms, but both can influence promoter melting via changes in DNA topology, thus explaining why they do not seem to have an additive effect (Figure 4B).

While at the full-length promoter Fis binding is sufficient to counter repression by H-NS (83), at the P1short promoter the KanR cassette plays a significant role at the sites found within H-NS bound regions, probably by both increasing RNAP binding to the promoter via an increase in negative supercoiling and by directly disrupting the H-NS bound region by its presence. This disruption is not as efficient at the *nsr3* position however, where the H-NS bound region is up to 15 Kb long and had been previously identified as a transcriptionally silent region (26).

The lack of an effect of the KanR cassette on the full-length P1long promoter underlines the role of Fis and H-NS to provide a constant local environment for *rrnB*P1 expression independently of the genetic neighbourhood. At the faster growth rates Fis binding already activates transcription enough so that the *rrnB*P1 promoter is no longer influenced by the local changes in supercoiling provided by the divergent promoter. At the slower growth rates, when Fis concentrations are lower and its activation effect smaller, repression by H-NS binding reduces the effect of the KanR cassette, or any other neighbouring gene effects (63).

### Local genomic context: H-NS

H-NS binding on the *E. coli* genome is not homogeneous but is often found in clusters of binding regions of different size (18). Extended H-NS binding regions are associated with lower levels of gene expression, often strengthened by its interaction with StpA (26, 27). H-NS binding is found more frequently at the ter half of the genome, due in part to the higher AT content. The P1short promoter used here is similar to the truncated version of the *rrnB*P1 promoter studied by Schroder and Wagner (63). In their case, the fragment was cleaved in the middle of the UP element, disrupting the second H-NS binding site. They observed a significantly weaker interaction of H-NS with this fragment compared to the full-length promoter (63). Therefore, H-NS binding should be decreased in the P1short promoter used here compared to the full-length promoter. A correlation between H-NS regions and gene expression is still observed, in agreement with previous work showing that strong repression within an H-NS rich tsEPOD can also occur for a promoter that is not specifically regulated by H-NS (5, 32).

H-NS binding regions overlap with five of the insertion sites studied with P1short, but not all of them are affected in the same manner, other parameters, such as the size of the H-NS-bound genomic regions and the level of the neighbour’s gene expression also play a role. One of the sites is within a tsEPOD (*nsr3*), two are next to highly expressed genes (*nsr2* and *left3)*, and two are not (*right1* and *ter4*). In the latter pair, the one that is found within the *ter* macrodomain is repressed more than the other is, as we had observed previously (31). At two of the positions studied here, *nsr2* and *left3*, the H-NS bound regions containing the reporter cassette are found downstream to a highly expressed gene. At these two positions, there is a significant decrease in expression of GFP. H-NS in its bridging DNA-binding mode can bind and stabilize the plectonemic structures that are formed by DNA supercoiling and can also block twist diffusion (84) creating a supercoiling domain boundary in the genome (85). In this case, the combination of the highly expressed gene towards the insertion site and H-NS binding could result in the stabilization of a structure particularly unfavourable to transcription initiation, as suggested by the low levels of expression of the genes found upstream of these insertion sites (SFig. 4A).

The results described here show that while the absolute values of the P1short promoter activity depend on genomic position due to local effects, of H-NS binding and neighbouring gene expression, the fold change as a function of growth rate remains the same when measured independently of gene copy number (SFig. 6). While the activity of H-NS and the amount of supercoiling can depend on local genomic context, the change in the activity of these same regulatory factors as a function of growth rate is independent of genome position. So for example, promoters that are found at a highly expressed position, such as *ori3* and *nsl3*, have the same growth rate dependence than those with a low level of expression such as at *nsr2*.

### Transcription interference

In addition to changes in DNA topology, convergent transcription can result in transcriptional interference (TI) when the RNA polymerase transcribing a neighbouring gene does not stop and continues on the convergently expressed gene, thus leading to a head-on clash of transcription complexes and the synthesis of antisense RNA sequences (2). A previous study has shown that TI appears to be widespread for convergent genes in the *E. coli* genome (1), due to inefficient transcription termination. The strength of the promoter can influence the result of the competition between the two convergent RNA polymerases by changing the frequency of initiation, the number of RNAP on the gene and thus the probability of reaching the end without encountering an RNAP coming from the opposite direction (86). Activation of the promoter by either Fis or the divergent cassette can thus further increase gene expression in the case where transcription interference is present, such as at the *nsr2* position. H-NS has been shown to act as a silencer of recently acquired DNA (87); however, transcription interference can also play a significant role in gene silencing, particularly when the insertion occurs within or next to a highly expressed gene (38). Alternatively, depending on gene orientation, it can result in the opposite effect, by integrating the newly acquired gene within an operon organisation (6).

### Conclusions

Other examples have been described of genes that are insulated by the upstream binding of Fis (88), and even by idle RNAP, showing that a divergent promoter does not necessarily need to be expressed to influence gene expression (89). Other abundant nucleoid proteins, such as HU, can have a strong effect on the changes in gene expression as a function of genome position (41), suggesting that this NAP can also act as an insulator of neighbouring gene transcription. The lack of HU in fact causes important overall changes in the organisation of the bacterial nucleoid (14, 22).

Here we have provided evidence for a further level of transcription regulation by nucleoid proteins and neighbouring gene orientation that plays an important role in maintaining the levels of gene expression independent of local effects. This can apply not only to the seven different ribosomal operons promoter regions but could have a more general role in the organisation at the supra-operon organisation level, given the large number of Fis and H-NS binding sites along the genome and specifically at the boundaries of the supra-operons (6).

Within the packed bacterial genome, a divergent gene orientation is found more often than a convergent orientation (3, 86). In light of the results presented here this suggests that this orientation could be favoured for maintaining the expression of the two genes dependent on each other, but less dependent on their local genomic environment. Insulation from neighbouring genes can contribute to a higher probability of success of horizontal gene transfer, define independently expressed genomic regions such as pathogenicity islands and help explain the effect of transcription on the formation of chromosomal interaction domain boundaries (90). Furthermore, modification of neighbouring genes’ effects, including the efficiency of termination and transcription interference, has been recently shown to play a role in the evolution of changes in gene expression levels during growth under pressure from antibiotics (91). Finally, knowledge of the different mechanisms that can insulate gene expression from its genomic neighbourhood can be used to build more stable synthetic genetic networks (92).

## Supporting information

SFig

## ACKNOWLEDGEMENTS

The authors would like to thank Aswin Seshasayee for the data on gene expression and David Lee for the pDoc-K plasmid.

