## Supplementary material for "Insulation of ribosomal promoter activity by Fis, H-NS or a divergent promoter within the packed *E. coli* genome": SFig

### Supplementary materials. Brambilla *et al*

A

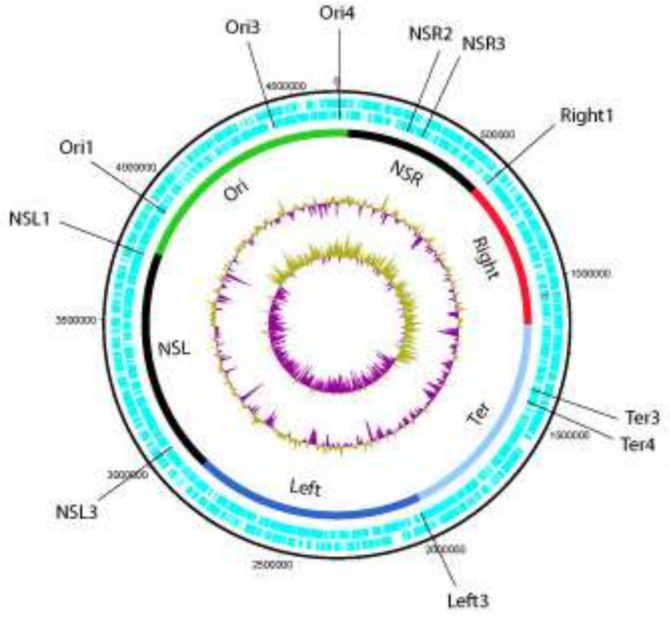


B

**P1**

-35 -10

TTGCGCGGTCAGAAAATTATTTTAAATTTCCTC<u>TTGTCA</u>GGCCGGAATAACTCCC<u>TATAAT</u>GCGCCACCACTGACA

16-bp spacer

**P5**

-35 -10

ACAACATCTAAGAGAAAAATTATA<u>TTGACA</u>TCTGCCCTTGAATAAGC<u>TATAAT</u>AGTAGTCTTAGTTAGAGAAGGAGGGTATAA

17-bp spacer

**Supplementary Figure 1. A. Position of the insertions in the genome.** The macrodomains as defined by Valens *et al.* (1) are shown, the %GC plot and the GC skew [(GC)/(G+C)]. Plot made with DNAPlotter (2). **B. Sequences of the *rrnB*P1 and P5 promoters.** The discriminator region is highlighted in red, the UP element in green and the transcription start site in grey.

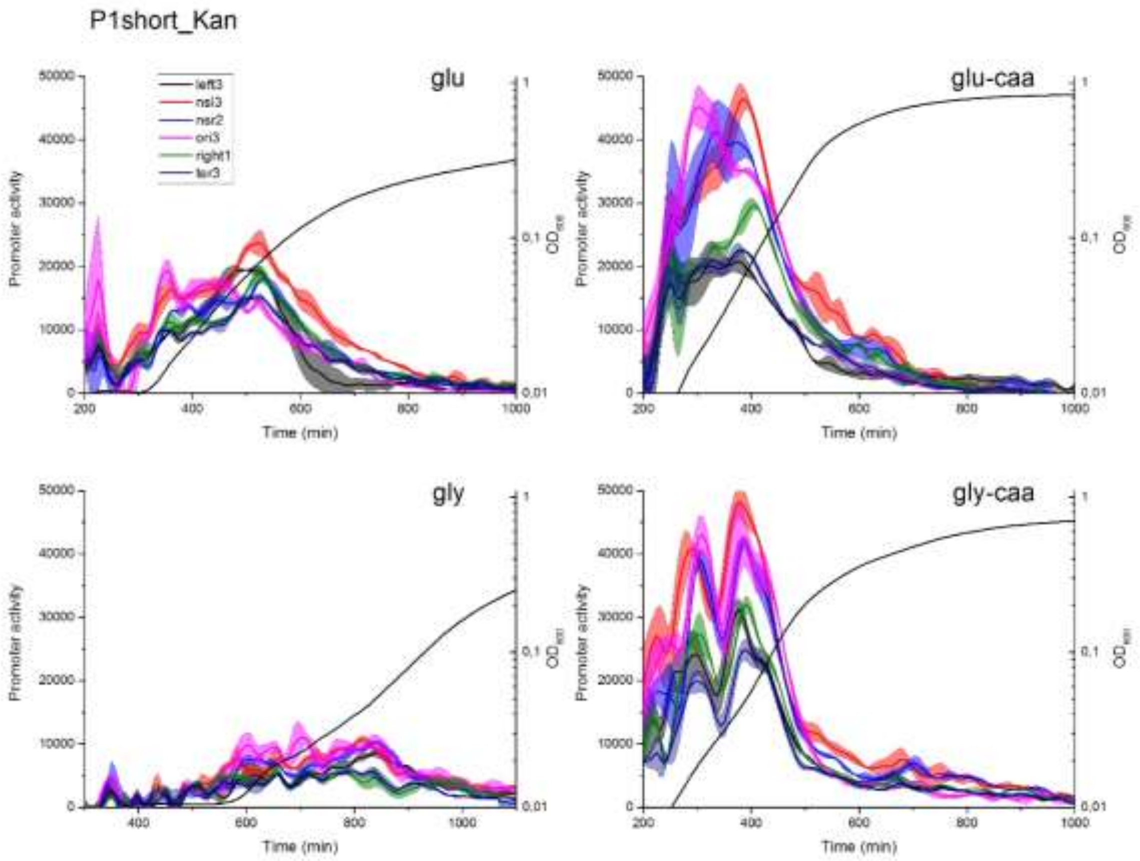


Supplementary Figure 2. Promoter activity and $OD_{600}$ for the reporter strains at five positions and four growth media.

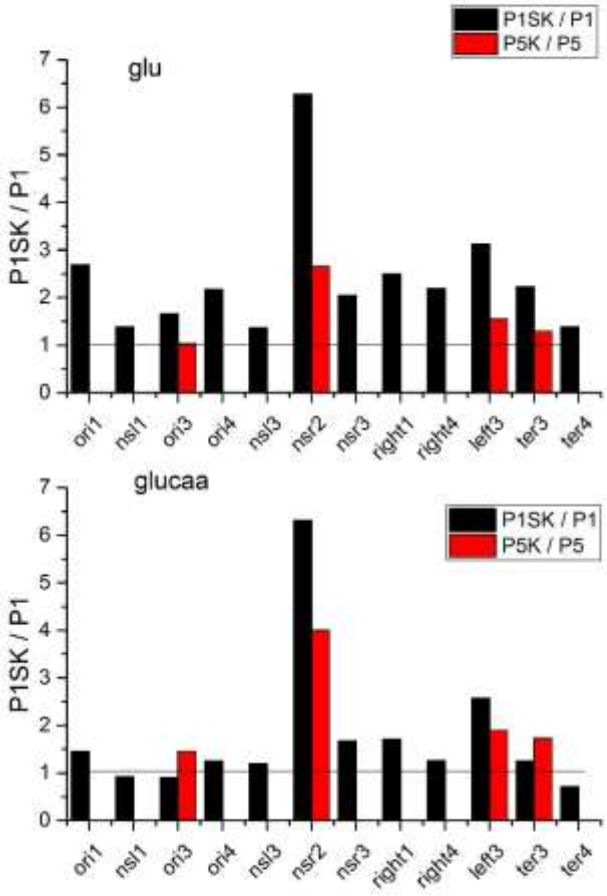


**Supplementary Figure 3.** The ratio of P1short in the presence and absence of the divergent KanR cassette, black bars, compared to the ratio of the activity of the P5 promoter in the presence and absence of the same cassette, red bars, in M9-glu, top panel and M9-glucaa, bottom panel.

A

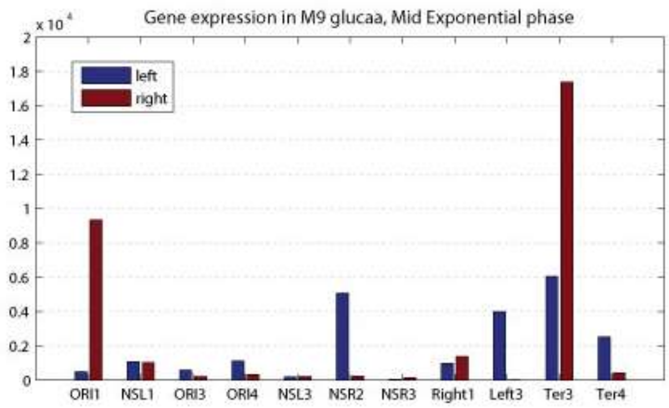


B

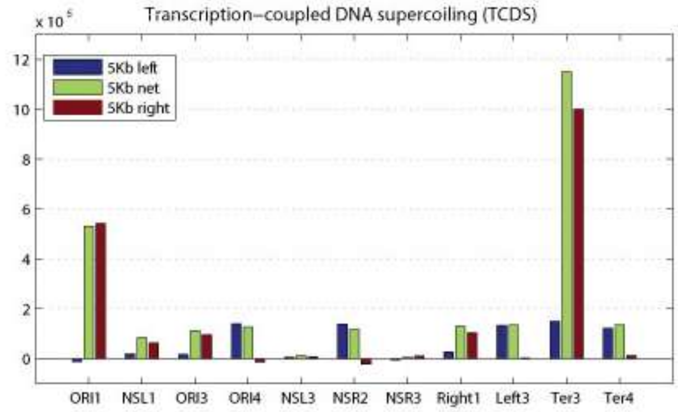


C

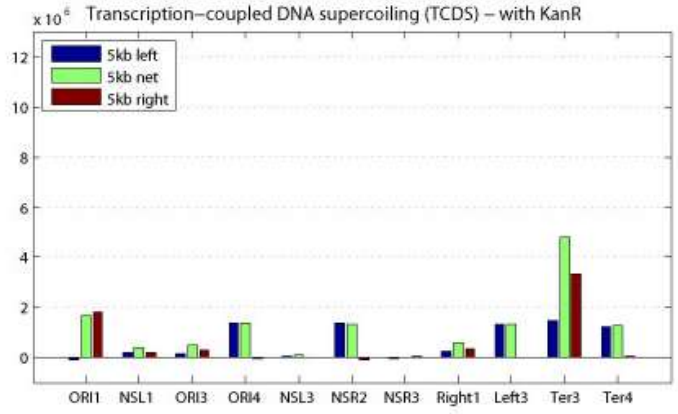


**Supplementary Figure 4**. A. Transcription level of the closest neighboring genes on the left and right of the insertion site in middle exponential phase in M9-glucaa (data from (3)). **B.** TCDS calculated as described by (4) from the transcription activity in a 5 Kb window on either side of the insertion site and the resulting, “net”, TCSD at the site. **C.** The same calculation carried out by adding 1.2 Kb on the right side of the insertion site, to mimic the presence of the KanR cassette.

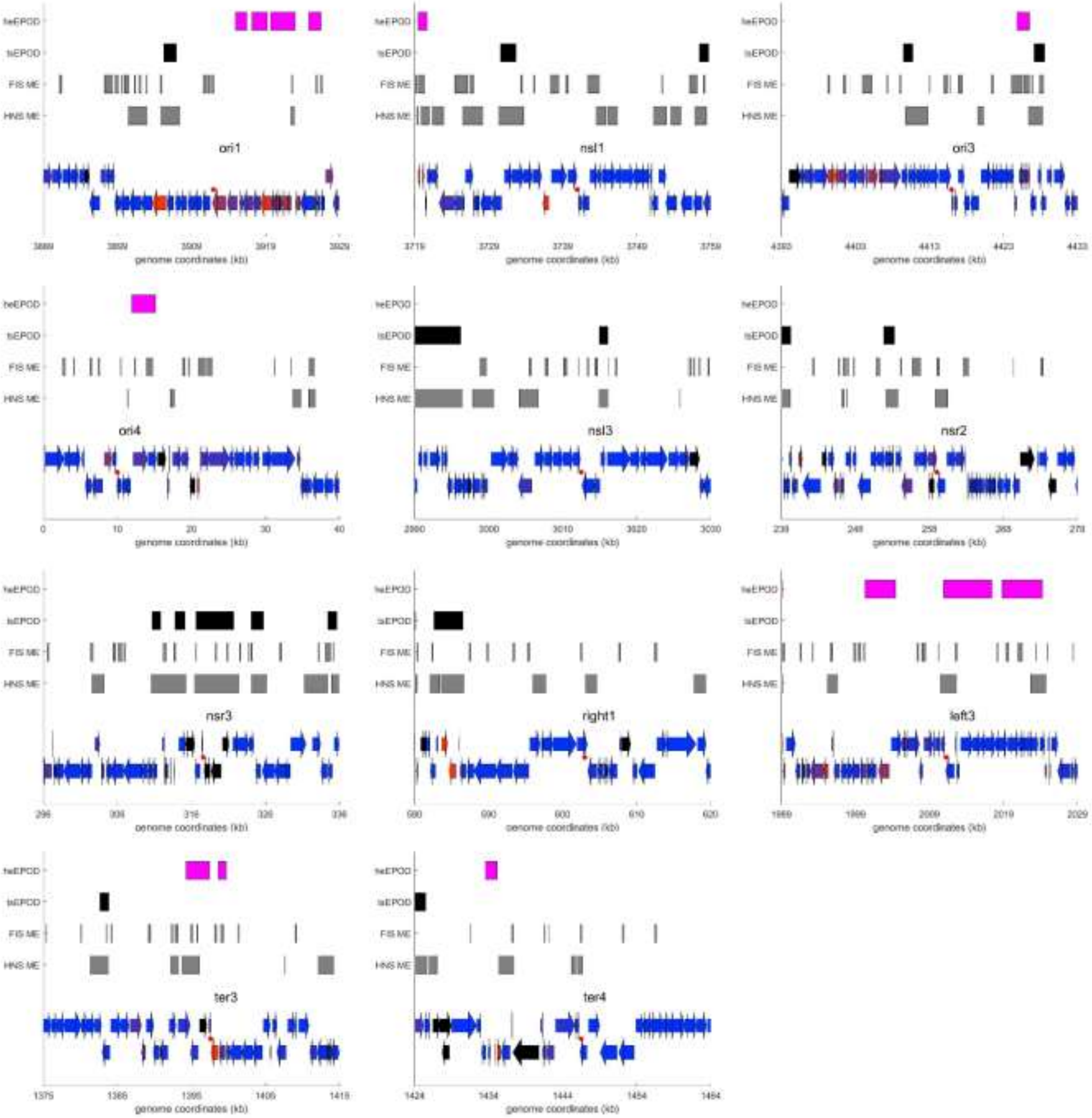


**Supplementary Figure 5.** For each insertion position are shown over a 40 Kb window: the neighboring genes, the binding of H-NS and Fis in mid exponential phase (ME) from (5) as well as the presence of Transcriptionally Silent Extended Protein Occupancy Domains (tsEPOD) and Highly Expressed Extended Protein Occupancy Domains (heEPOD) from (6). The gene color corresponds to the number of reads from blue to red, with blue equals 0 and red set at 1500 reads from the dataset from (3) in M9-glucaa in mid-exponential phase. The black genes did not have expression data.

A

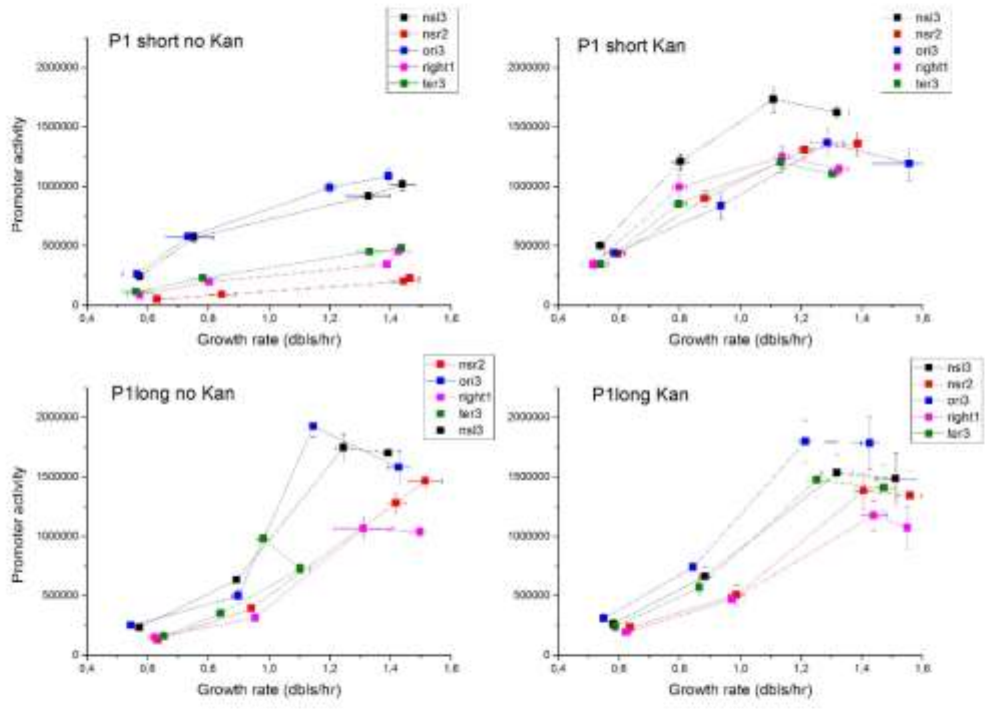


B

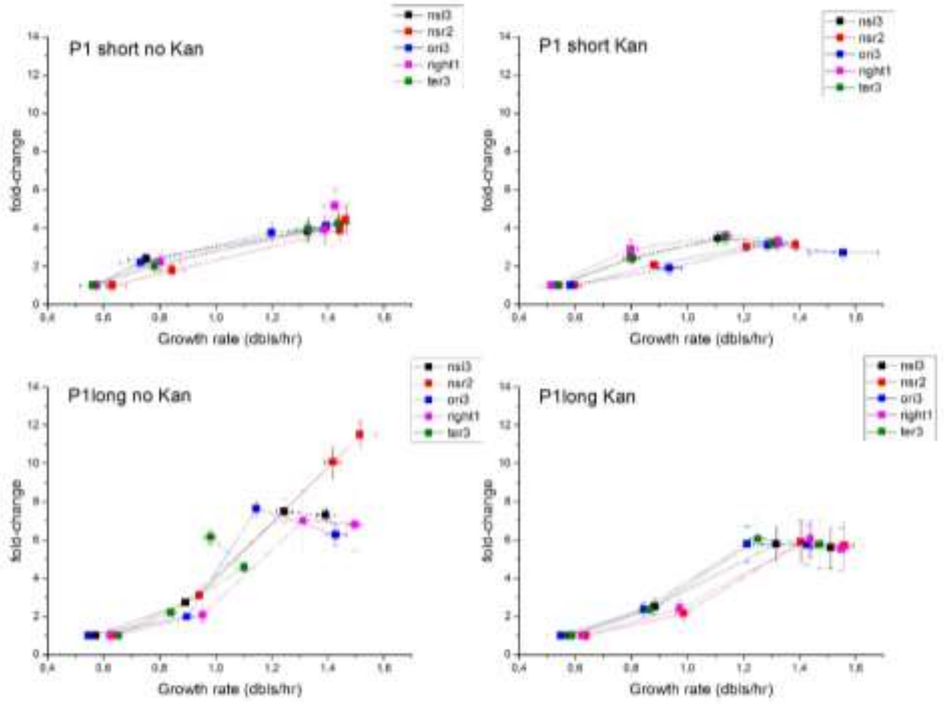


**Supplementary Figure 6. A**. Growth rate dependence of promoter activity after normalization for gene copy number at five positions in four growth media, M9-gly, M9-glu, M9-glycaa and M9-glucaa. **B.** Fold-change in promoter activity as a function of growth rate relative to the promoter activity in M9-gly.

1. Valens,M., Penaud,S., Rossignol,M., Cornet,F. and Boccard,F. (2004) Macrodomain organization of the Escherichia coli chromosome. *EMBO J*, **23**, 4330–41.

2. Carver,T., Thomson,N., Bleasby,A., Berriman,M. and Parkhill,J. (2009) DNAPlotter: circular and linear interactive genome visualization. *Bioinformatics*, **25**, 119–120.

3. Lal,A., Krishna,S. and Seshasayee,A.S.N. (2016) Regulation of global transcription in E. coli by Rsd and 6S RNA. *bioRxiv*, 10.1101/058339.

4. Sobetzko,P. (2016) Transcription-coupled DNA supercoiling dictates the chromosomal arrangement of bacterial genes. *Nucleic Acids Res.*, **44**, 1514–1524.

5. Kahramanoglou,C., Seshasayee,A.S.N., Prieto,A.I., Ibberson,D., Schmidt,S., Zimmermann,J., Benes,V., Fraser,G.M. and Luscombe,N.M. (2011) Direct and indirect effects of H-NS and Fis on global gene expression control in Escherichia coli. *Nucleic Acids Res.*, **39**, 2073–2091.

6. Vora,T., Hottes,A.K. and Tavazoie,S. (2009) Protein occupancy landscape of a bacterial genome. *Mol. Cell*, **35**, 247–253.
